# Application of RGB trichrome staining to the study of human parasites

**DOI:** 10.1101/2022.09.20.508650

**Authors:** Juan Diego González-Luna, Francisco Gaytán

**Affiliations:** Department of Cell Biology, Physiology and Immunology, Faculty of Medicine and Nursing, University of Córdoba, Spain; Instituto Maimónides de Investigación Biomédica de Córdoba, Córdoba, Spain

## Abstract

Human parasitic infections are major contributors to global disease load, compromising the human life and resulting in considerable morbidity and mortality worldwide. However, many parasitic diseases have been neglected and little investigated in western medicine. Although recently developed molecular techniques have revolutionized the taxonomy of parasites and the parasitic disease diagnosis, histopathology still remains a powerfull tool for the analysis of parasitic diseases, allowing direct observation of the parasite thus providing information about the morphological features of the parasite ifself, and revealing tissue alterations at the parasite-host interface. The recently developed RGB trichrome (acronym for the primary dye components, picrosirius Red, fast Green, and alcian Blue) stains the main components of the extracellular matrix, specifically collagens and proteoglycans. We have applied the RGB tricrome staining to human tissues infected by the main three classes of human parasites: Protozoa (*Leishmania donovanii* and *Toxoplasma gondii*), helminths (*Trichinella spiralis, Enterobius vermicularis, Dirofilaria* spp. and *Echinococcus granulosus*) and ectoparasites (*Demodex folliculorum* and *Demodex brevis*). Trichrome stain results in detailed staining of the parasite microanatomical structure, and highlights host tissue alterations such as granulomatous inflammation, immune cell infiltrate, or increased amount of collagen as a sign of parasite-induced fibrosis. Yet, the use of RGB trichrome, as a complement of hematoxylin and eosin staining, provides additional valuable information to assess parasitic infection histopathology.

## Introduction

Parasitic infections constitute a major concern about public health worldwide. It has been estimated that more than 3 billion people are infected by parasites^**1–4**^. In general, infections are more frequent in developing countries, mostly at the tropical areas, and several parasitic diseases have been listed as neglected tropical diseases by the World Health Organization (WHO)^**5**^. However, parasites have spread all over the world with human migrations, and currently there are many emerging, and re-emerging, parasitic diseases out of their classical endemic areas due to the rise in migration and international travels, together with climate change and the increase in the number of immunocompromised patients; furthermore, poly-parasite infections seem to be more frequent than previously suspected^**6,7**^.

The number of parasite species is difficult to ascertain. An indirect method is to consider how many parasite species affects every free-living known host, and infer the ratio of parasites to free-living species. Different studies have proposed different proportions ranging from 5% to more than 50%, but parasitic species probably outnumber free-living species^**8–11**^. However, the specificity of each parasite with respect to the host is not well known for many species. Anyway, it is apparent that the parasite lifestyle is ubiquitous, and the role of parasites in ecosystems and in human evolution has been systematically underestimated. Yet, it has been scarcely considered in the literature the possible role of parasites preventing or modifying some diseases, either by competing with other harm parasites or modulating and setting up host immune system.

Infectious diseases are produced by viruses, bacteria, protozoa, fungi, helminths, and to a lesser extent, arthropoda. Between these, only Protozoa, Helminths and Ectoparasites (arthropods) are strictly considered as parasites^**1**^. The medical and veterinary importance of parasites is enormous. Depending on the study, about 70 protozoan and 250 helminth species have been described as the causative agents of human parasitic diseases, some of which are zoonotic diseases transmited by animals^**2,12**^ that also have a high impact in livestock species. Many of these parasitic infections affect preferentially people from undeveloped countries and research on this topic has been neglected to a greater or lesser extent in developed contries^**5**^. However, the emerging prevalence of parasitic infections suggests that parasitology should receive more attention in general biological research as well as in medical education.

The diagnosis of parasite infections have been classically established by clinical data and morphological criteria by direct observation of tissue samples under the microscope^**13**^. Currently, the application of molecular-based techniques has modified the diagnostic process, as well as the classification and taxonomy of parasite species^**12,14,15**^, as in many cases morphological characteristics do not allow classification beyond the genus level, and new genotypes, some of them re-classified now as new species, have been recognized by applying new molecular and immunohistochemical techniques^**12**^.

Even so, histopathological observation of infected tissue samples still provide valuable information. First, the direct visualization of the parasites is a unequivocal evidence of the parasitic infection, and their morphological characteristics, such as size and shape, may allow its classification, at least up to the genus level. Second, the interface between the parasite and host tissues provide valuable data about tissue damage, inflammatory reaction, and the precise anatomical location of the parasite. This makes that careful histopathological examination of infected tissues or cytologic smears still constitutes an important tool that add valuable information to molecular-based procedures. Although hematoxylin and eosin (H&E) is the basic stain for microscopic obbservation, different staining methods such as trichrome stains are used for certain purposes. Thus, Masson trichrome stain has been used for identification of helminth eggs or pratozoan parasites in stool samples^**16,17**^. The observation of the host-tissue interface could reveal the existence of immune cell infiltration and reactive fibrosis in inflammatory areas. In this context, we have applied a recently developed trichrome stain^**18**^, the RGB trichrome (RGB is the acronym for the three primary dyes used: sirius Red, fast Green and alcian Blue). The sirius red and alcian blue, that stain specifically collagen and glycosaminoglycans/proteoglycans respectively, together with a general background protein stain (fast green) provide a wide spectrum of staining colors, that could be valuable for the study of the parasite structure itself, and the characteristics of the parasite-host interface. The objective of this study was to assess the usefullness of RGB trichrome for the staining of different human parasites in tissue sections.

## Materials and Methods

Human tissue samples were obtained from the Bio-archive of the Department of Pathology of the Faculty of Medicine and Nursing. These specimens were obtained for diagnostic purposes to confim suspected parasitic infections or after surgery due to different pathologies. Skin biopsies were also used for the study of demodex mites in the areas of the biopsy free of pathology. In all cases, sections of those samples in which the presence of parasites was reported in their pathology report were used for RGB staining. Additional appendix samples were studied for the presence of food residuals that mimicked parasites. These samples were collected between 1980-2000, in keeping with contemporary legislation, including informed consent. Anyway, the use of these samples was approved by responsible members of the Department of Pathology, granting patient confidentiality and following strict adhesion to current legislation.

Human tissues had been fixed in 4% phosphate buffered formaldehyde and embedded in paraffin. The number of samples for each parasite class were: 2 Protozoa (1 *Leishmania donovanii;* 1 *Toxoplasma gondii*), 16 helminths (2 *Trichinella spiralis*, 7 *Enterobius vermicularis*, 1 *Dirofilaria* spp. and 6 *Echinococcus granulosus*), 18 Ectoparasites (10 *Demodex folliculorum*, 8 *Demodex brevis*). In addition, some tissue samples indicating the presence of fungi were also used. Six μm-thick sections were cut from paraffin blocks, and after dewaxing and rehydration were submitted to RGB staining, following previously described methods^**18**^. Briefly, the sections were sequentially stained with 1% (w/v) alcian blue in 3% aqueous acetic acid solution (pH 2.5) for 20 min, washed in tap water, stained with 0.04% (w/v) fast green in distilled water for 20 min, washed in tap water, and finally stained with 0.1% (w/v) sirius red in saturated aqueous solution of picric acid for 30 min, washed in two changes of acidified (1% acetic acid) tap water, dehydrated in 100% ethanol, cleared in xylene, and mounted with synthetic resin. Some sections from the files of the Department of Pathology, that were already stained with hematoxylin and eosin, were also used. All figures in this study were stained with RGB trichrome unless otherwise indicated in the figure legend. A scale bar has been added in at least one panel per figure.

## Results and Discussion

The three main classes of parasites responsible for human diseases are Protozoa, Helminths and Ectoparasites, that show (in this order) an increasing structural complexity. Microscopic observation of infected tissues is a useful tool for the identification of parasites and the evaluation of tissue damage. We have investigated the usefulness of RGB trichrome for the staining of tissues infected with some prevalent parasites belonging to these classes.

### Protozoa

Protozoa are the infectious agents of a large number of severe human diseases that remain a major public health concern worldwide^**1,2**^. Two examples of protozoan infections, a skin lesion caused by *Leishmania donovanii*, and a placental section showing a tissue cyst of *Toxoplasma gondii* (one of the most successful human parasites)^**19**^ are shown in Figure 1.

**Figure 1.**
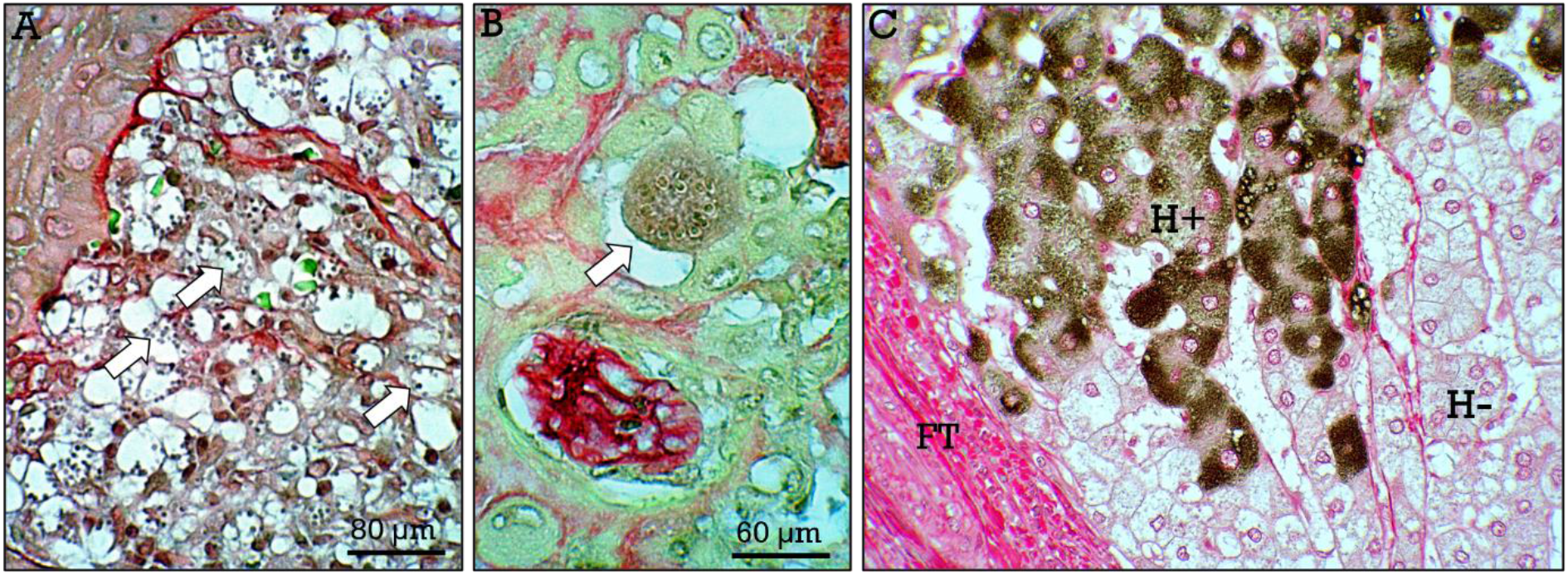
*Protozoa and RGB counterstaining*. Tissue sections from skin (**A**) infected with *Leishmania donovanii*, showing parasites inside macrophages (***arrows***) and from placenta (**B**) showing a tissue cyst (***arrow***) containing bradyzoites of *Toxoplasma gondii*. In **C**, RGB trichrome has been applied as background counterstain to a section of liver immunostained for HBsAg with the peroxidase method. Positive (***H+***) and negative (***H -***) hepatocytes and fibrous tissue (***FT***) can be observed.

As protozoa are unicellular organisms and the RGB trichrome is not particularly a cellular stain, the usefullness of this method is limited when applied to protozoan parasites and does not add valuable information to H&E stained tissues with respect to the structure of the parasite itself. However, as a complementary stain, RGB trichrome could provide valuable information about host tissue status and parasite-related alterations in the extracellular matrix. Furthermore, a potential advantage of RGB trichrome is its use as counterstain when applying diaminobenzidine (DAB)-peroxidase immunohistochemistry for the detection or identification of parasites. As an example, from a non-strictly parasitic infection, a liver section immunostained for the hepatitis B virus surface antigen (HbsAg) counterstained with RGB trichrome is shown in Fig. 1C. It can be appreciated that trichrome stain does not interfere with the DAB-derived chromogen, and allows concomitant detection of possible alterations in infected tissues, such as parasite-induced fibrosis.

### Helminths

#### Nematodes

Intestinal helminth infections (including Nematodes, Cestodes and Trematodes) affect more than 2 billion people around the world, being more prevalent in developing countries^**3,5**^. Nematodes (roundworms) are complex animals, that present nervous, gastrointestinal, excretory, muscular, and reproductive systems, with an internal pseudocoelomic cavity and an external collagenous cuticle^**20**^. There are a huge number of free-living and parasitic nematode species, and this phylum is extremely relevant in biomedical sciences, not only for being the causal agent of many parasitic infections, but because a free-living nematode, *Caenorharbditis elegans*, is one of the most used model in biomedical research, particularly in genetics and developmental biology^**21**^. We have studied three parasitic species of nematodes, *Trichinella spiralis, Enterobius vermicularis*, and *Dirofilaria* spp to assess their characteristics after staining with RGB trichrome. The biology and epidemiology of these species have been widely considered in the literature^**22–24**^. In this study we will make a brief description of the parasite life cycle and biology, focusing on the staining characteristics after application of the RGB trichrome.

##### Trichinella spiralis

*T. spiralis* is the causative agent of human trichinosis, a re-emerging infection that constitutes a serious zoonotic disease worldwide. *T. spiralis* has a broad range of host species and is an unusual parasite that completes its life cycle (comprising two generations alternating enteric and skeletal muscle stages) in the same host, and develops intracellularly during the larval (muscular) stage^**22**^. Briefly, upon ingestion of the encysted larvae from infected muscle meat, they are released, invade the intestinal mucosa, mature into adult worms, mate and reproduce. The newborn larvae released by gravid females, migrate through the lymphatic and blood vessels to the striated muscle, where they penetrate individual muscle cells and are encysted creating an unique intracellular habitat. This is a complex process in which one larva invades a striated muscle cell, and acts (in a virus-like pathway) modifying the striated muscle cell by controlling gene expression and inducing modifications in the infected cell and surrounding tissue, such as degradation of cytoskeleton, induction of the sinthesis of specific proteoglycans, and the formation of an enveloping collagenous capsule and new blood capillary vessels, thus turning it into a nurse cell, where the encysted larva is nourished and protected against host defense mechanisms^**25,26**^. In RGB stained sections of infected muscles (Fig. 2), nurse cells that are considerably enlarged with respect to non-infected muscle cells, showed bluish stained cytoplasm, and are surrounded by a red-stained collagen capsule (Fig. 2A,B). Several cross-sections of the larva appears in the cytoplasm. In general, larvae are green stained whith the exception of stichocytes, that are unicellular glands in the gastrointestinal system, that are red stained (Fig. 2C,D). Transverse cuticular ridges can be clearly appreciated in logitudinal worm sections (Fig. 2G). Frequently, nurse cells are surrounded by an inflammatory reaction containing eosinophils (that appears green-stained). The bluish staining of the remaining cytoplasm of nurse cells is likely due to the increased amounts of proteoglycans, as the synthesis of some of them (i.e., glypican-1) is induced in the nurse cells by the parasite^**25**^, together with the absence of the muscle cell cytoskeleton (that stain green) that has been degraded in nurse cells. The vascular network surrounding nurse cells can be also observed (Fig. 2H). In some nurse cells larvae are not present (Fig. 2I) but there are abundant cells inside the cyst. These likely correspond to nurse cells in which the larva has died and has been invaded by inflammatory cells. Fibrosis is evidenced by increased amounts of red-stained collagen surrounding muscle cells (Fig. 2H,I).

**Figure 2.**
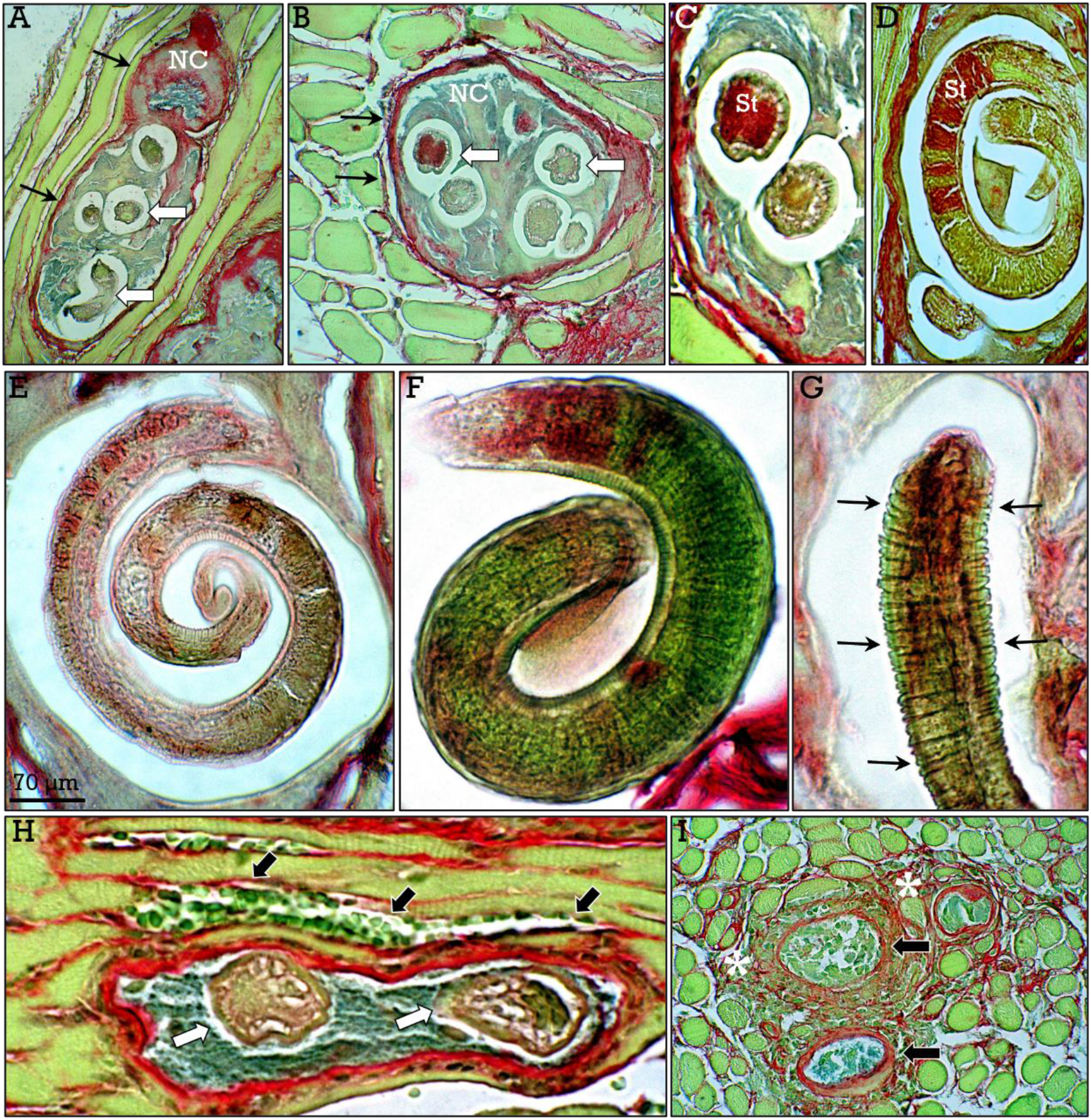
*Nematodes I. Trichinella spiralis*. In infected muscles, nurse cells (***NC***) can be observed in longitudinal (**A**) and transverse (**B**) sections, containing transected trichinella larvae (***white arrows***) and encased by a collagenous capsule (***black arrows***). Stichocytes (***St***) are shown in transverse (**C**) and longitudinal (**D**) sections. Longitudinal sections showing the spirally coiled larva (**E,F**) and their transverse cuticular ridgets (***arrows*** in Blood vessels are abundant around nurse cells (***black arrows*** in **H**). Degenerating nurse cells (***arrows*** in **I**) showing infiltration by inflammatory cells and surrounding collagenous tissue (***asterisks***).

Microscopic observation of the larvae of *T. spiralis* in affected muscles can be performed satisfactorily by routine staining with H&E. Hoewever, some advantages of RGB trichrome staining, specially for preclinical studies on trichinellosis, are an adequate observation of the encysted larvae and, at the same time, highlighting the potential fibrosis in the affected areas by the specific red staining of collagen, that can be objectively quantified under polarized light microscopy^**27**^.

##### Enterobius vermicularis

*Enterobius vermicularis* (pinworm) is a human pathogen with a high prevalence worldwide, affecting to more than a billion people, preferently children, and occasionally adults^**23**^. The human seems to be the only host of *E. vermicularis*, and the infection has a harmless course in most cases. Pinworms have a simple direct cycle in humans. Upon ingestion of eggs the larvae hatch, mature, and are established in the large intestine, mainly in the cecum and appendix. At night, gravid females migrate to the anus and lay eggs in the anal folds, where skin-adhered eggs produce pruritus and induce scratching. In this way, contamination of hands and subsequent invasion of the digestive system maintains the chain of infection through self-inoculation.

In this study we have assessed the staining properties of *E. vemicularis* present in appendectomy pieces. Pinworms were found in the appendix lumen, and were easily recognizable by their morphological features including their longitudinal alae (Fig. 3), and organs consisting of a collagenous cuticle, peripheral muscular system, pseudocoelom, gonads (testis or ovaries) and abundant eggs inside gravid females (Fig. 3G-J). The cuticle stains redish (Fig. 3B) while the worm stains basically green, although some component of the digestive system, such as the pharinx bulb displays a mixture of colors (Fig. 3E). In gravid females, the egg cuticle shows a green-stained outer layer and a red-stained inner layer, encasing an embryo that stains green (Fig. 3I,J). Although in most cases there was a low number (2-5) of pinworms per appendix, high number of parasites were found occasionally (Fig. 3F). Usually, the infection is mild and causes only discomfort in children, being even asymptomatic in adults. However, several studies have suggested that the presence of pinworms in the appendix may lead to acute appendicitis^**28,29**^, raising the possibility that *E. vermicularis* infections could be more serious than previously suspected.

**Figure 3.**
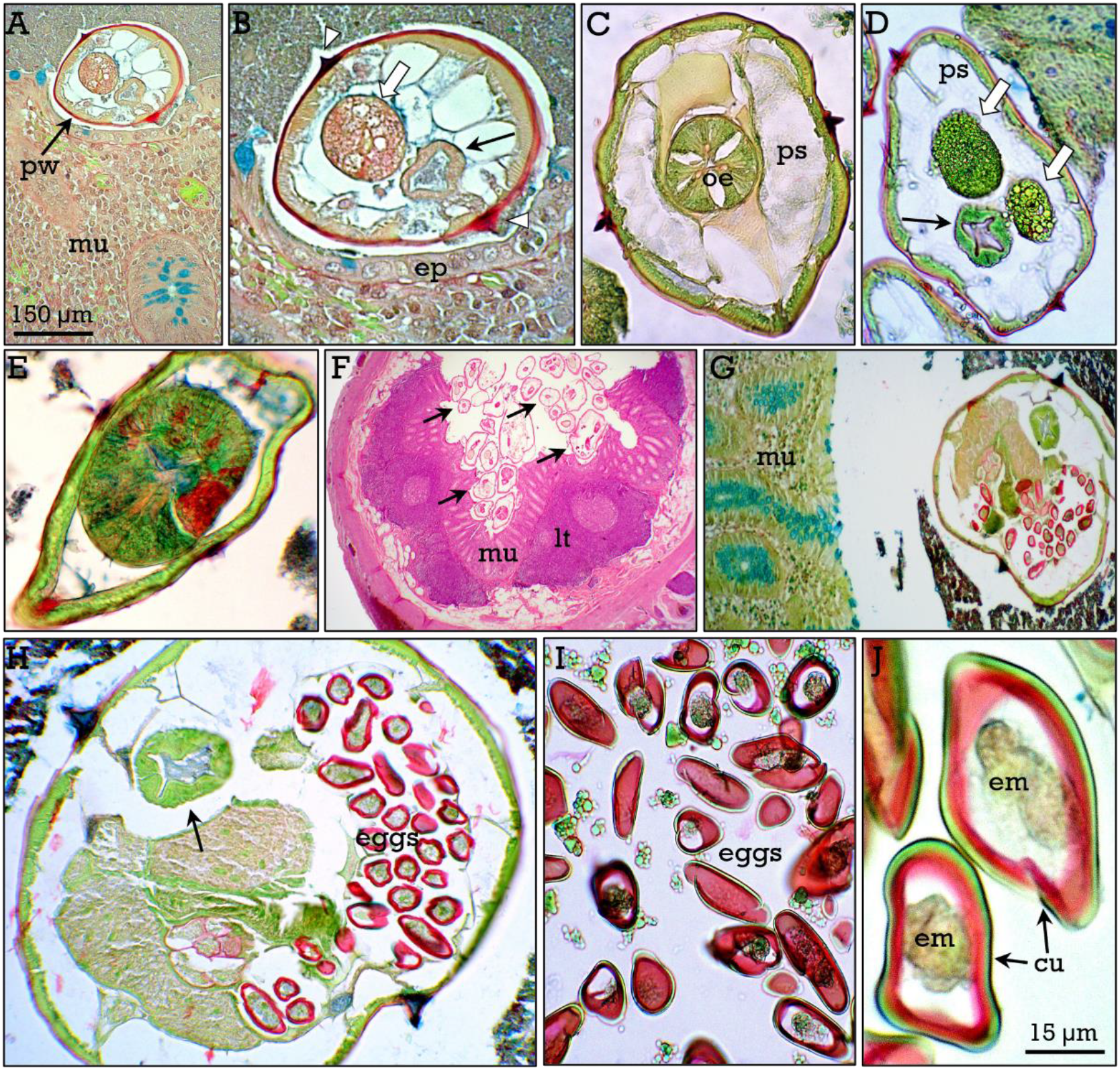
*Nematodes II. Enterobius vermicularis*. Pinworms were found in the lumen of the appendix, and the epithelium (***ep***) and the lamina propria of the mucosa (***mu***) were apparently intact. Figures **A-E** show sections of *E. vermicularis* showing the red-stained cuticle, cuticular alae (***arrowheads***), pseudocoeloma (***ps***), digestive system (***black arrows***), triradiate oesophagus (***oe***), pharingeal bulb (**E**) and gonads (***white arrows***). In some samples, large numbers of pinworms were found in the lumen of the appendix (**F**). Figures **G,H** show a gravid female (***arrow*** in **G**) containing abundant eggs, that show a cuticle (***cu***) with an external green-stained, and an internal red-stained (**I,J**) layer, encasing the embryo (***em***). **F**, H&E

Staining with RGB trichrome provides clear contrast between the different tissues and internal organs of the worms. Notably, the brilliant staining of the eggs of *E. vermicularis*, raises the possibility that RGB trichrome staining could be a useful tool for the identification of eggs from different nematode and/or other helminth species, and deserves comparative studies for this purpose.

While studying the appendix samples to assess *E. vermicularis* staining, we noticed the presence of occasional undigested seeds and plant residuals that can be eventually confused with parasites and, in fact, several studies have addressed this issue^**30–33**^. In this context, we decided to investigate the staining outcomes of seeds and plant tissue remnants after application of RGB trichrome as a method that contributes to its differentiation from parasites. Figure 4 shows seeds and different plant-derived structures that were found in the appendix, that mymicked parasites, particularly when these animals were distorted due to death or to supervening alterations during tissue processing (Fig. 4H). The results indicate that RGB trichrome results in a particularly brilliant staining of seeds and plant-derived tissues, thus facilitating its differentiation from parasites or parasite-derived structures.

**Figure 4.**
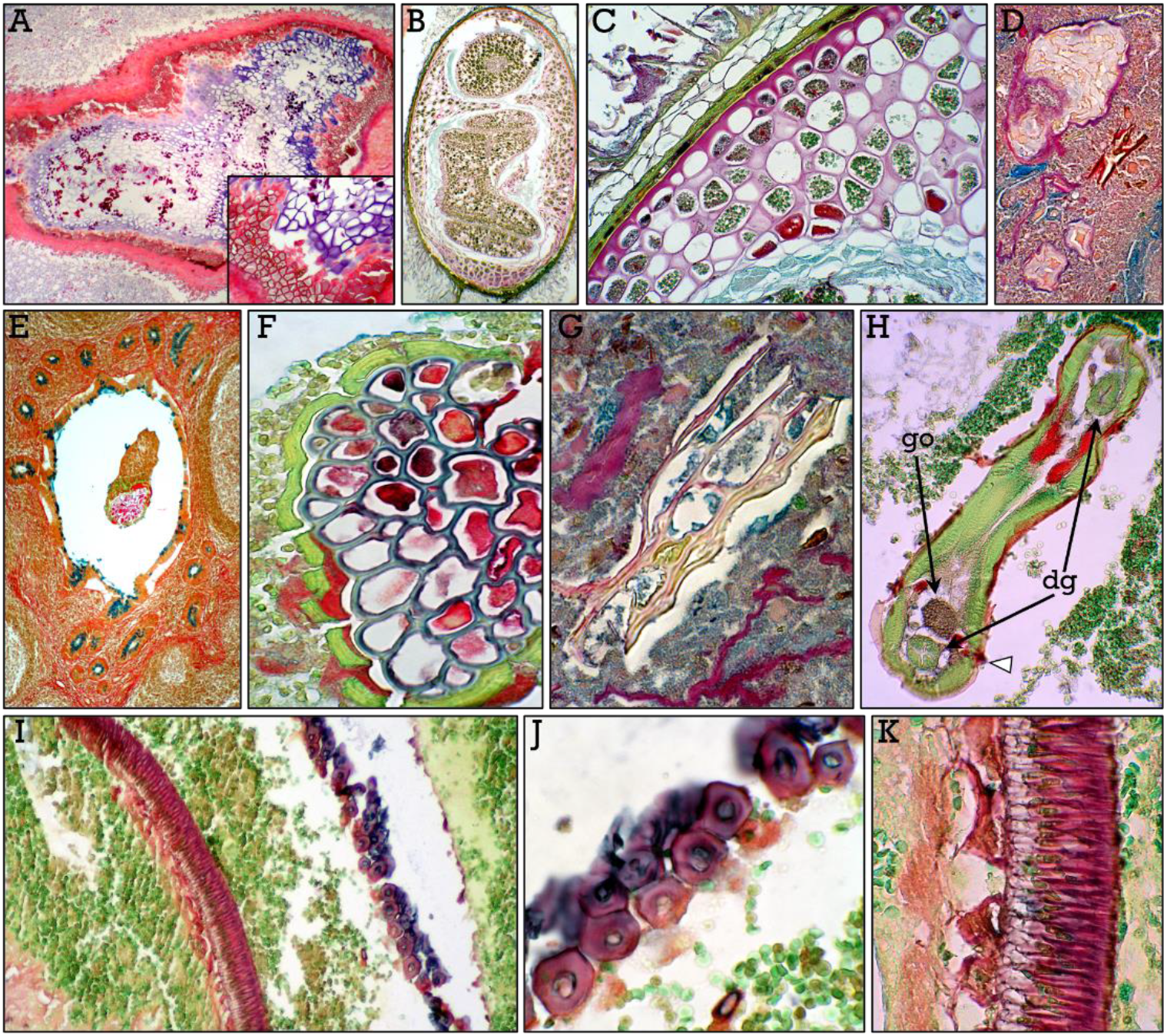
*Plant remnants in the appendix*. Undigested vegetal materials that could mimic parasites found in the appendix. Micrographs show plant remnants (**A,E,F**), a seed (**B,C**), starch granules (**D**), and sclereids (**I-K**). Plant structures show a thick, intensely stained cell wall (**A,E,F**). Some vegetal structures (**G, K**) can be particularly confusing when worms are degraded or distorted by tissue processing (**H**). Distinctive features (such as alae in *E vermicularis; **arrowheads*** in **H**), as well as digestive system (***dg***) and gonads (***go***) allows parasite identification.

##### *Dirofilaria* spp

Different species of the genus *Dirofilaria*, such as *Dirofilaria repens*, and *Dirofilaria immitis*, are the causative agents of dirofilariosis, a zoonotic parasitic infection transmited by different mosquito species, and whose prevalence is increasing worldwide. Dirofilarias parasite canid and felid species and the main reservoir for human infections are dogs, and less frequently cats^**34**^. Adult worms produce microfilariae that are present in the peripheral blood from which they are ingested by mosquitoes, thus creating the infection chain. Although dirofilariosis is a neglected parasitic disease considered as sporadic, its actual incidence constitutes a global health problem^**24,34,35**^.

Humans are incidental hosts, in which usually the parasites do not reach sexual maturity^**36**^. Infections by *Dirofilaria repens* produce local inflammatory nodules that can be found in virtually any anatomical location^**24**^, and could be confused with lesions of tumoral origin. In this study, the nematode was found in a nodule in the submucosa of the small intestine, encapsulated by a local granulomatous inflammatory reaction. Crossections of the worm showed a red-stained cuticle containing longitudinal ridges, a prominent subcuticular layer of muscle cells, as well as a digestive system lined by a layer of cells with apical microvilli surrounded by the pseudocoeloma (Fig. 5).

**Figure 5.**
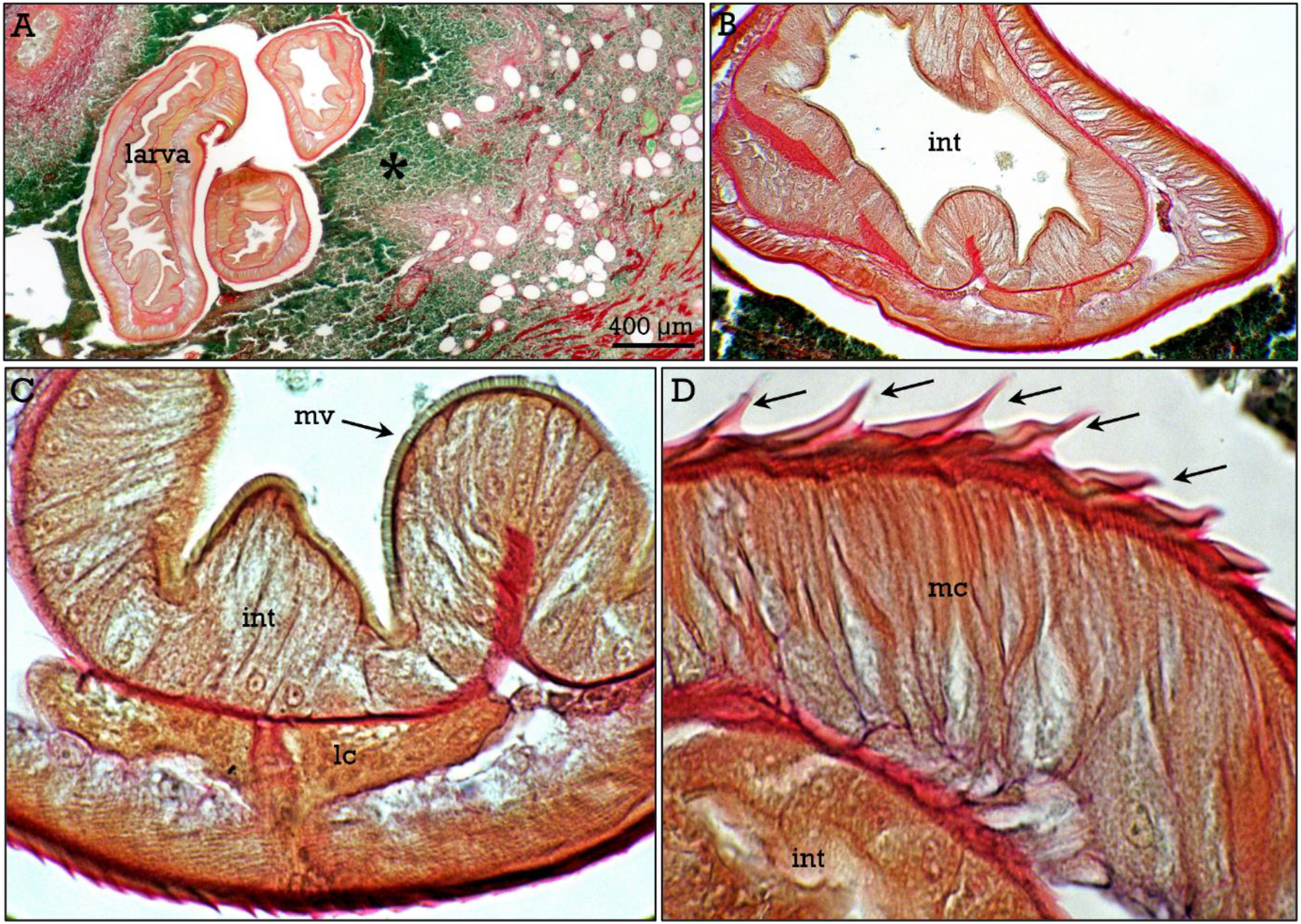
*Nematodes III. Dirofilaria spp*. Sections of the worm located in the intestinal submucosa (**A**), surrounded by a granulomatous inflammation (***asterisk***). The intestine microvilli (***mv***), longitudinal cuticular ridges (***arrows***), muscular cells (***mc***) and lateral chord (***lc***) can be observed (**B-D**).

The morphology of the worm, including the longitudinal cuticular ridges, and the general anatomic features strongly suggest that they correspond to *Dirofilaria repens*, that is the most frequent species involved in human dirofilariosis^**24**^; however, as these morphological features are not exclusive of this species, we have considered it as an unidentified *Dirofilaria* spp.

The host response to parasitic infection involves local inflammation at the parasite-host tissue interface. In accordance, the worm found in the intestinal submucosa was surrounded by a granulomatous inflammatory reaction, and an intense infiltration of green-stained eosinophil leukocytes (Fig. 6), that constitutes the hallmark of the host anti-helminth inflammatory response^**37**^. Huge numbers of eosinophils infiltrate both the intestinal lamina propria and the submucosa (Fig. 6), and coarse collagen fiber bundles were abundant in the infiltrated areas of the submucosa, likely reflecting the initial steps of parasite-induced fibrosis.

**Figure 6.**
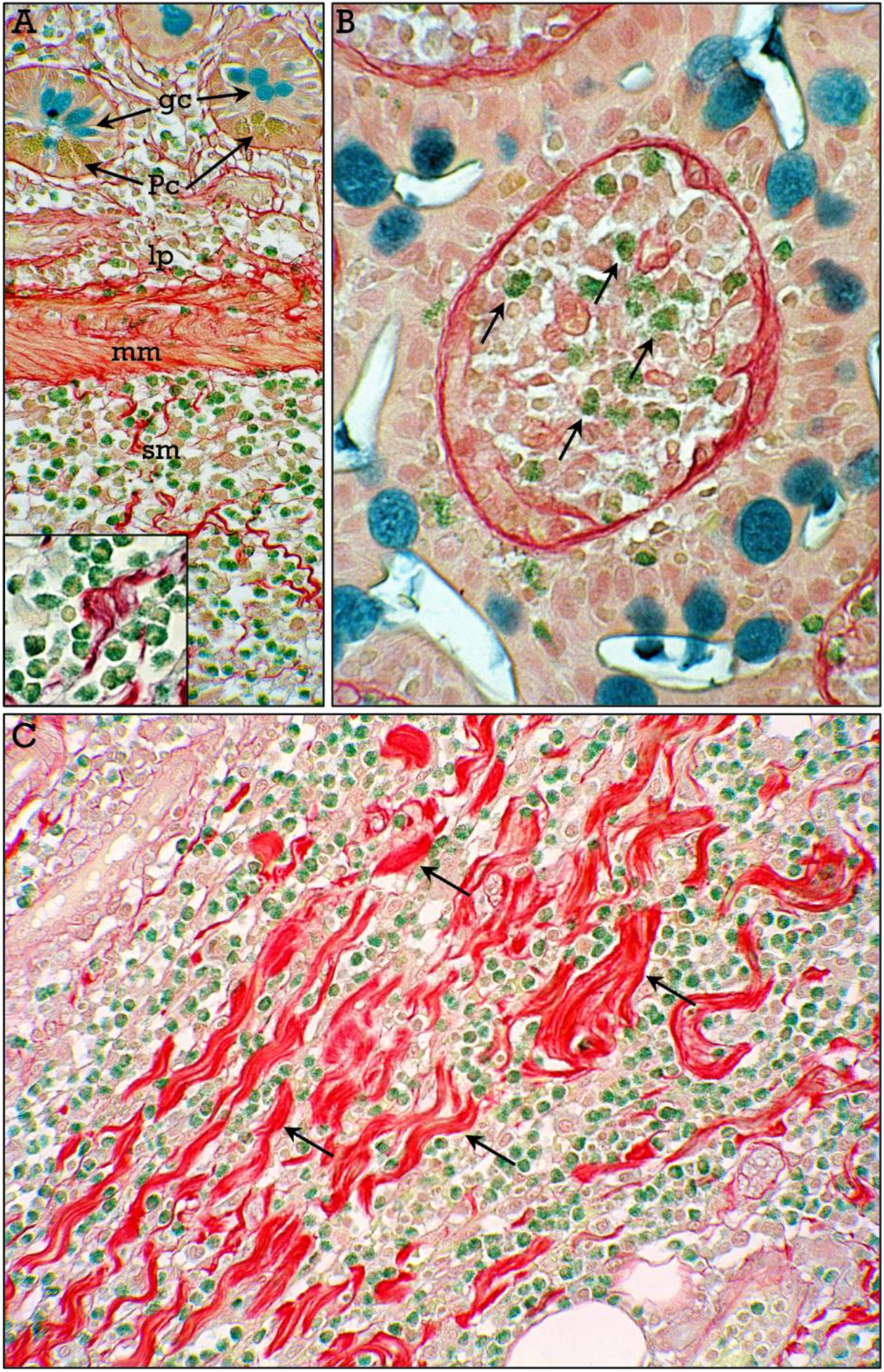
*Nematodes IV. Eosinophil leukocytes infltration*. Surrounding the dirofilaria larva, abundant eosinophil leukocytes (***arrows*** in the ***inset***) infiltrate the lamina propria (***lp***) of the intestinal villi (***arrows*** in **B**), and the submucosa (***sm***) in which coarse collagen bundles (***arrows*** in **C**) can be observed. ***mm***, muscularis mucosae. ***Pc***, Paneth cells; ***Gc***, Globet cells.

#### Cestodes

##### Echinococcus granulosus

Cestodes (tapeworms) are helminths that causes serious infectious diseases. Echinococcosis is a zoonotic infection that is one of the 17 neglected tropical diseases recognized by the World Health Organization (WHO), and is caused by tapeworms of the genus *Echinococcus^**38**^*. The terminology applied to this genus and even to the structural characteristics of the larval stage in the intermediate host is rather complicated and a great effort have been made to reach a consensus for the terminology of the different aspects of this parasitic disease^**39**^. Cystic echinococcosis (CE; aka cystic hydatid disease) is caused by the larval stage (metacestode) of *Echinococcus granulosus*. This scenario is complicated because in this genus there are several genotypes (G1-G10), that vary in their biology, rate of development, pathogenicity, and geographical extent, and some of which are now considered as new different species. So, all genotypes causing CE are denominated as *E. granulosus sensu lato*, whereas genotypes G1-G3 are denominated *E. granulosus sensu strictu*^**40–42**^. As the most frequent human infections are due to genotypes G1-G3, and specially G1, the cases analyzed in this study likely correspond to *E. granulosus sensu strictu*. However, for the sake of simplicity, we are going to refer to the causative agent of the CE as *E. granulosus*. The life cycle of these cestodes implicates two different hosts. Briefly, mature tapeworms in the intestine of carnivorous, definitive, hosts (for instance dogs, that are the most important reservoir for human infection) produce eggs that are eliminated along with feces and thereafter ingested by herbivores, intermediate, hosts (sheeps or other livestock animals in human infection). In the intermediate hosts (human are accidental intermediate hosts) the ingested eggs develop into oncospheres that invade the intestinal mucosa and enter the circulatory system. Although the oncospheres can be established in virtually any organ, the most frequently affected are liver and less frequently lungs. In these organs, the oncospheres transform into a bladder-like structure denominated metacestode that accumulates fluid, and generally grows slowly and is initially asymptomatic. Clinical cystic echinococcosis may appear as a consequence of pressure on surrounding tissues due to cyst growth and/or to rupture of the cyst.

RGB-stained histological sections of metacestodes in human tissues are shown in Fig. 7. A total of 6 samples (4 from liver, 1 from lung and 1 from ovary cysts) were studied. The metacestode (i.e., cyst) is formed by an internal narrow germinal layer (cellular) and an external acellular chitinous layer, the laminated layer (Fig. 7A,B), that show a wide range of colors. These two layers derive from the parasite. Brood capsules are formed by budding from the germinal layer toward the cavity and contains the infective form of the larvae, known as protoscolex (Fig. 7C,D), that can be also found free in the cyst cavity. Surrounding the laminated layers there is the adventitial layer from host origin, and that in RGB stained sections appears in most cases as a double layer, an inner zone composed granuloma-like inflammatory cells, with occasional multinucleated cells (Fig. 8D) and an external layer of red-stained collagenous tissue (Fig. 7 and 8). The protoscoleces show a rostellum with two rows of green-stained hooklets (Fig. 7C-F) and abundant calcareous bodies (that could be blue- or brown-stained; Fig. 7D). Protoscoleces can adopt two morphologies, either invaginated or, occasionally, evaginated (Fig. 7E). Each protoscolex could give rise to an adult worm in the definitive host if ingested, or to new metacestodes in the intermediary host after metacestode rupture and spread of the cyst content (secondary hydatidosis), that can also cause anaphylactic shock. Some metacestodes are not fertile (ie., they lack protoscolex formation) and could be collapsed (Fig. 8B,C) or even calcified. In these case, the presence of laminated layer remnants and/or isolated hooklets (Fig. 8B,C) are hallmarks of cystic echinococcosis. Infiltrating eosinophil leukocytes were found in high numbers in the liver tissue around the cyst (Fig. 8E,F).

**Figure 7.**
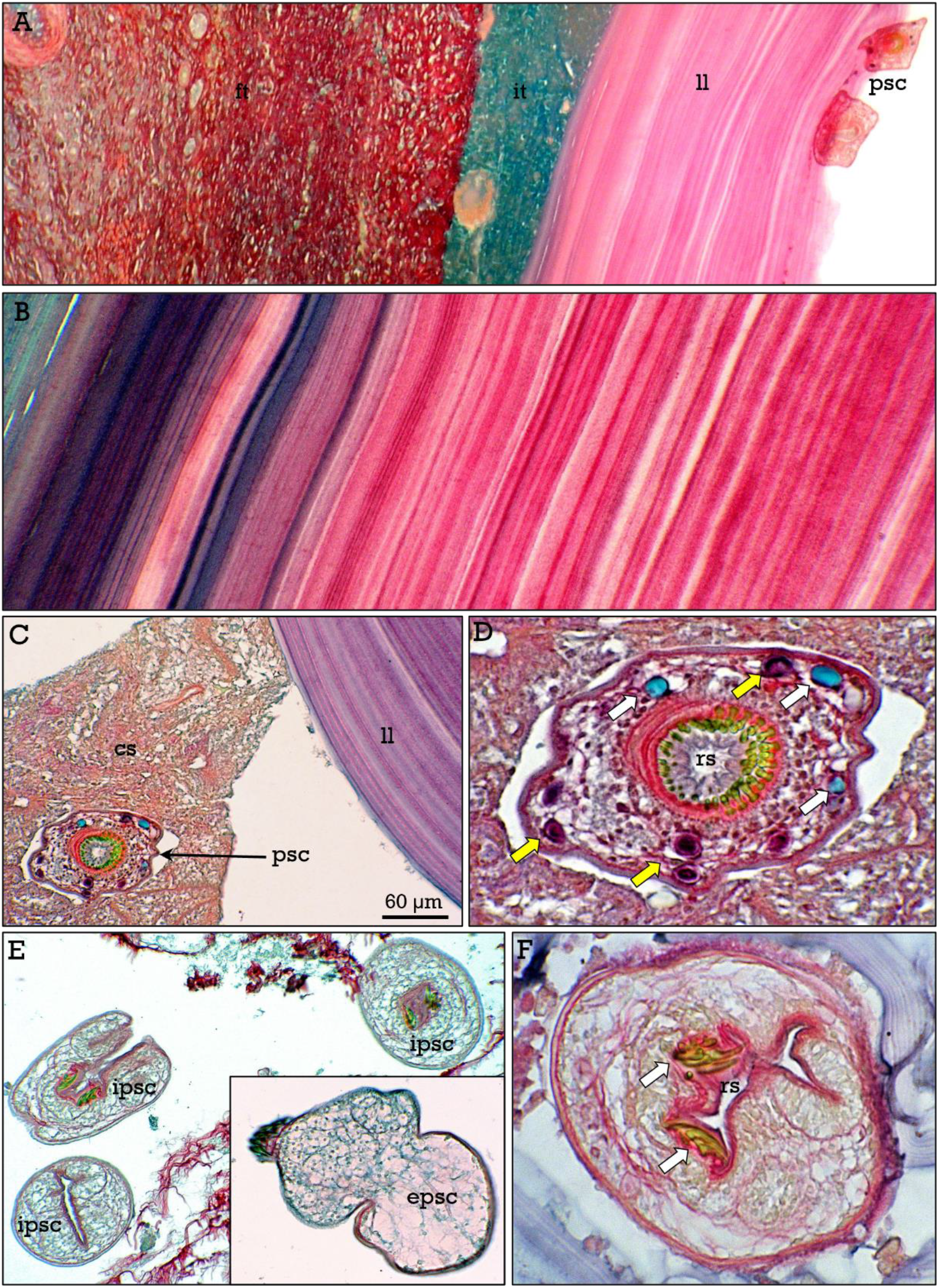
*Echinococcus granulosus I*. Cyst (**A**) showing the laminated layer (***ll***) and two displaced protoscoleces (***psc***) present in the lumen cyst, surrounded by inflammatory (***it***) and fibrous (***ft***) tissue. The multilaminated layer is shown at higher magnification in **B**. A protoscolex (***psc*** in **C**) and cyst sand (***cs***), are shown at higher magnification in **D**, showing the rostellum (***rs***) with the double row of hootlets and two types (blue- and brown-stained; ***white*** and ***yellow arrows***) of calcareous granules. Invaginated (***ipsc***) and evaginated (***epsc***) protoscoleces can be found (**E**). In **F**, a higher magnification of a protoscolex showing the hootlets (***arrows***) in the rostellum (***rs***).

**Figure 8.**
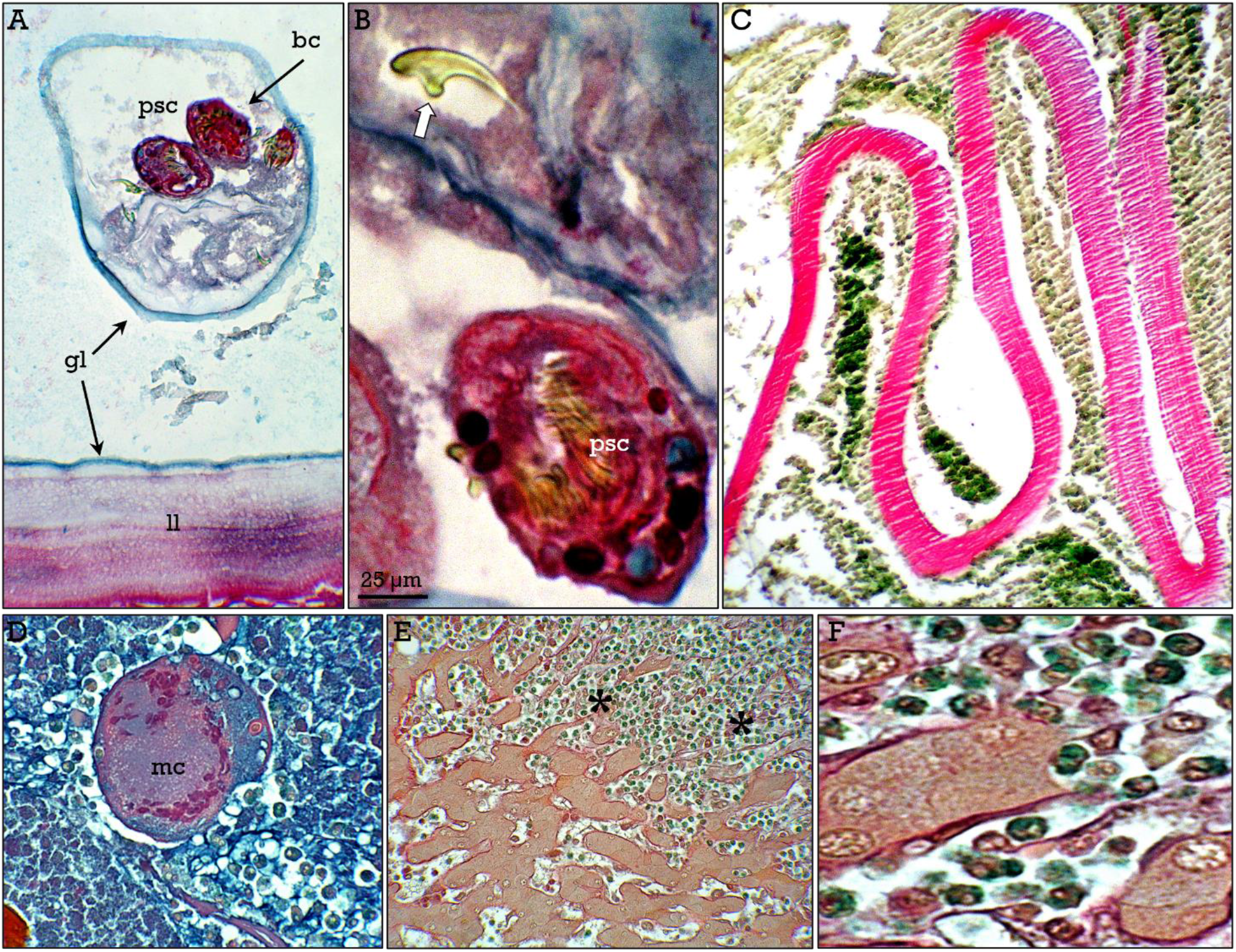
*Echinococcus granulosus II*. **A**, brood capsule (***bc***) surrounded by the germinal layer (***gl***) containing protoscoleces (***psc***). In **B**, one isolated hootlet (***arrow***) can be observed. In **C**, laminated layer in a collapsed cyst. Multinucleated cells (***mc*** in are occasionally present in the inflammatory tissue surrounding the parasitic laminated layer, and eosinophils are abundant (***asterisks***) in the surrounding liver tissue (**E,F**).

### Ectoparasites

#### Demodex

Demodex mites are microscopic arachnids resident in the skin belonging to the phylum Artropoda, family Acaridae, and are the most complex components of the skin microbiota. There is a large number of species-specific mites. In humans, two mite species inhabit the pilosebaceous units: *Demodex folliculorum* located at the hair follicle infundibulum, and *Demodex brevis* located deeper in the sebaceous glands^**43–45**^. Humans are the only host of these two species and they are possibly present in almost every adult person, being (if this is so) the most frequent and successful of the human parasites. It is thought that demodex mites feed on dead skin cells and sebum. However, as these mites cannot be kept alive in vitro, there are many uncertities with respect to their biology, physiology, and the mechanisms of their possible pathogenic effects. We are going to describe the two species separately, since there are significant differences in both their habitat and morphology.

##### Demodex foliculorum

*D. folliculorum* are found in the infundibulum of the hair follicles, and are particularly abundant in the face, although they can be present in hair follicles of any anatomical location. Their body structure, characteristic of the Acaridae family, consist of a gnathosoma (containing the mouthparts) and the podosoma (containing four pairs of legs) that correspond together to the cephalothorax, and the opisthosoma, that corresponds to the abdomen. In spite of their complexity, they are microscopic animals measuring from 280 to 300 μm, depending on the sex. The general morphology and the structural details than can be appreciated after RGB staining are shown in Figs. 9 and 10. The cuticle stain red and show characteristic transversal striations in the opisthosoma that facilitate the identification of mites even when only part of the opisthosoma is present in the section (Fig. 9A,F). The distal part of the opisthosoma is rounded (Fig 9A,C,F), which is a distinctive feature of this species, with respect to the *D. brevis*. In the gnathosoma some parts of the oral structure can be apreciated. In the podosome, the legs, leg-related muscles, salivary glands, and the nervous synganglion can be appreciated in adequate longitudinal (Fig. 9) and tranverse (Fig. 10) sections. Eggs (greenish-stained) are arrow-shaped in longitudinal sections and can be observed inside the opisthosome of female mites (Fig 9C), or free in the hair follicle infundibulum (Fig. 10F) after egg laying. Frequently, several mites can be found in the same follicle (Figs. 9D, 10A-D), sometimes closely accommodated, difficulting individual identification.

**Figure 9.**
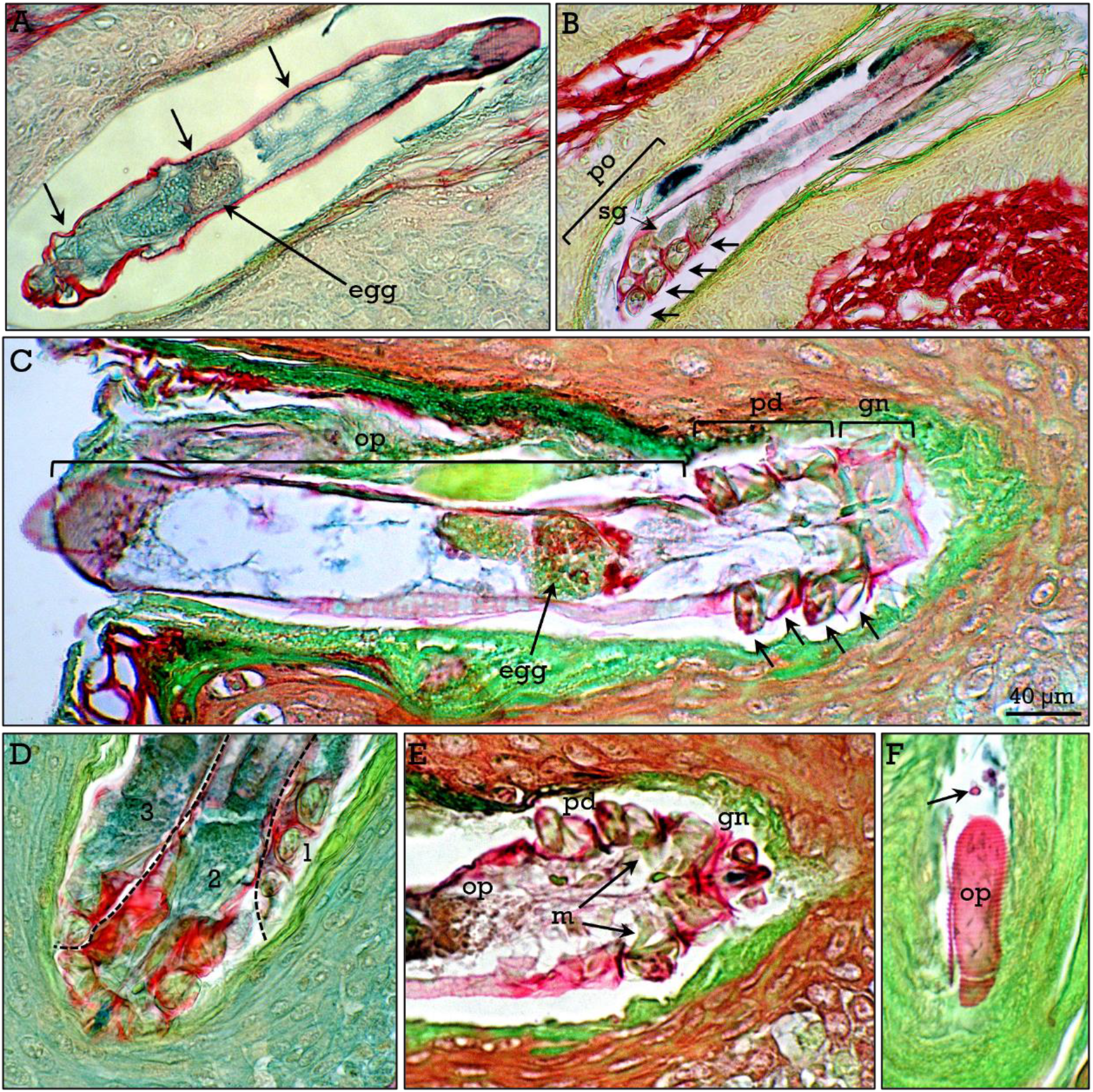
*Demodex folliculorum I*. Longitudinal sections (**A-C**) of *D. folliculorum* in the hair infundibulum showing different mite structures such as the cuticle (***large arrows***), gnathosoma (***gn***), podosoma (***po***), opisthosoma (***op***), legs (***short arrows***), synganglion (***sy***), salivary glands (***sg***), muscles (***m***) and eggs. In **D**, three mites (delineated by dashed lines) are closely accommodated in a hair follicle. In **F**, tangential section of the opisthosoma (***op***) of a mite in the hair infundibulum. ***Arrow***, fungus.

**Figure 10.**
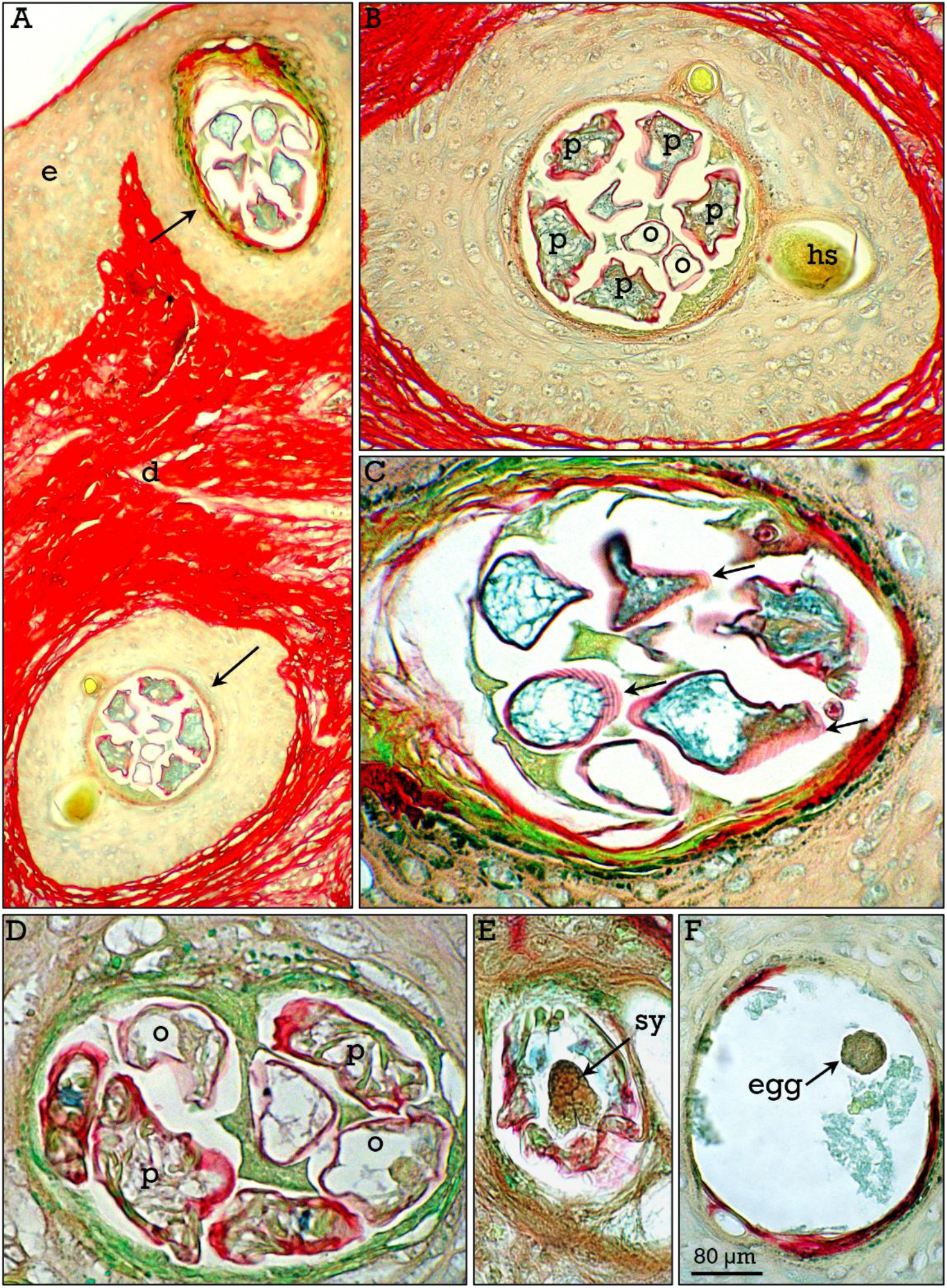
*Demodex folliculorum II*. Transverse sections of several *D. folliculorum (**arrows***) in hair infundibula sectioned at podosoma (***p***) or opisthosoma (***o***) level (**A-D**). The synganglion (***sy*** in **E**) and a released egg (**F**) in the infundibulum can be observed.

##### Demodex brevis

*D. brevis* are found deeper in the pilosebaceous units, located in the sebaceous glands. They are slightly shorter than *D. folliculorum*, measuring from 150 to 220 μm. Their structure (Fig. 11) is close similar to that of *D. folliculorum*, except for the opisthosoma tip that is sharp instead of rounded (Fig 11C), and eggs that are oval in longitudinal sections. The different shape of the opisthosoma allows discrimination between both species, together with their different locations in the pilosebaceous unit. However, the anatomical location is not always a discriminative sign. For instance, although *D. brevis* are usually located at the sebaceous glands, it cannot be discarded that could be also found in the hair follicle infundibulum, since this is the only presumable way for the mites to move from a sebaceous gland to another. Otherwise, the distinction between both species is rather difficult in transverse sections if anatomical location is unclear. *D. brevis* were frequently found in pairs (Fig. 11G), but also in groups, particularly in the sebaceous glands of the nipple (Fig. 11A). The different anatomical structures such as legs (Fig. 11B), synganglion (Fig. 11D), and eggs either inside (Fig. E) or outside (Fig. 11F) the mite can be observed. Specimens likely corresponding to immature protonymphae (Fig. 11H) showing six legs and a shorter opisthosoma were occasionally found.

**Figure 11.**
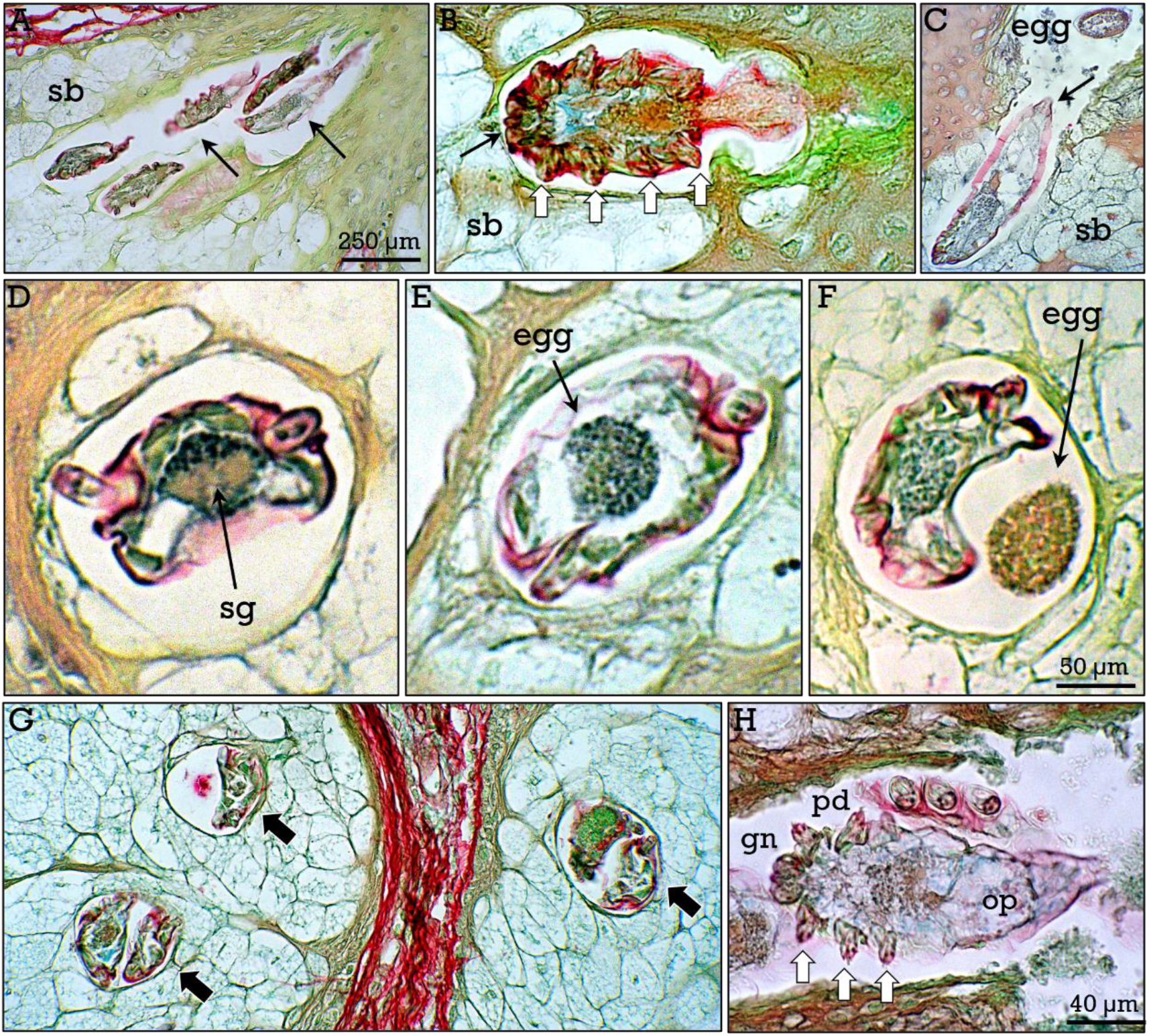
*Demodex brevis*. Longitudinal (**A-C**) and transverse (**D-G**) sections of *D. brevis*. Several mites can be found in the same sebaceous gland (***arrows*** in **A, G**). The gnathosoma (***arrow***) and the legs (***white arrows***) in the podosoma are shown in **B**. The characteristic pointed end of the opisthosoma (***arrow***) can be observed in **C**. Several mite structures such as synganglion (***sg***), and eggs inside (**E**) and outside (**F**) the mite can be observed in **D-F**. In **H**, a *D. brevis* showing the morphological features of an immature protonympha: six legs (***arrows***) and a short opisthosoma.

In spite of the existence of numerous studies reporting an association between the density of demodex and some skin inflammatory reactions such as acne rosacea and marginal blepharitis^**46,47**^, definitive data beyond statistical association has not been reported and the role of demodex in these human skin processes remain controversial, particularly because the mechanisms underlying these alterations are poorly understood. In fact, demodecosis (i.e., alterations due to demodex mites) are difficult to differentiate from idiopathic skin inflammatory diseases. It is therefore unclear if in humans the mites behave as comensals without deletereous effects. In this study we have found some histological signs suggestive of the implication of mites in skin inflammatory processes, as the presence of granulomas containing mite remnants (Fig. 12A-D), in which was not possible to ascertain to what species did they belong. Furthermore, some sebaceous glands showed an apparent degradation of cells (Fig. 12 E,F) together with the presence *D. brevis*. Anyway, a causal relationship between the presence of mites and these alterations cannot be ascertained from these data.

**Figure 12.**
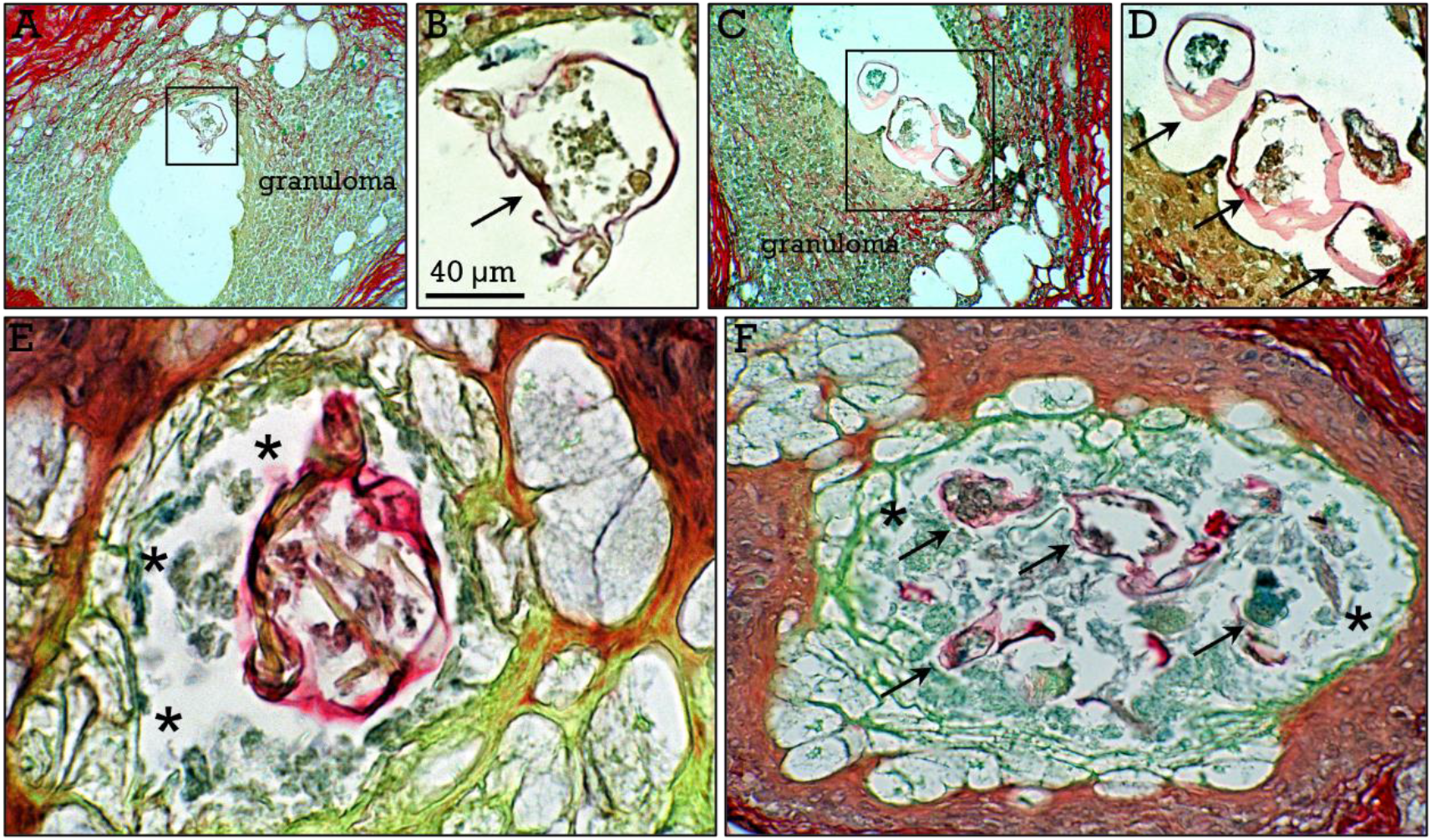
*Skin alterations related to demodex mites*. Granulomas (**A,C**) containing demodex remnants (***arrows***), recognizable at higher magnification (**B,D**). In **E,F**, apparent degradation of cells in sebaceous glands (***asterisks***) containing a mite (**E**) or mite remnants (***arrows*** in **F**).

The relationships between *Demodex* mites and skin bacteria are also unclear. Yet, while mites have associated bacteria, it has been proposed that mites can also represent a defense against pathogenic bacteria by releasing free fatty acid from the triclycerids in the sebum through their lipase enzyme activity^**48**^. In this study we have noticed that in some skin samples showing abundant granular blue-stained bacterial colonies in the hair follicles (Fig. 13A,B), accumulation of blue-stained material (presumably bacteria) was also found inside *Demodex* mites, that can be appreciated in either logitudinal (Fig. 13C) or transverse (Fig. 13D,E) mite sections. This raises the possibility that *Demodex* feed on bacteria in the hair infundibulum, and opens up a series of possibilities regarding the interaction between mites and bacteria and their possible roles either modulating or preventing skin bacterial infections.

**Figure 13.**
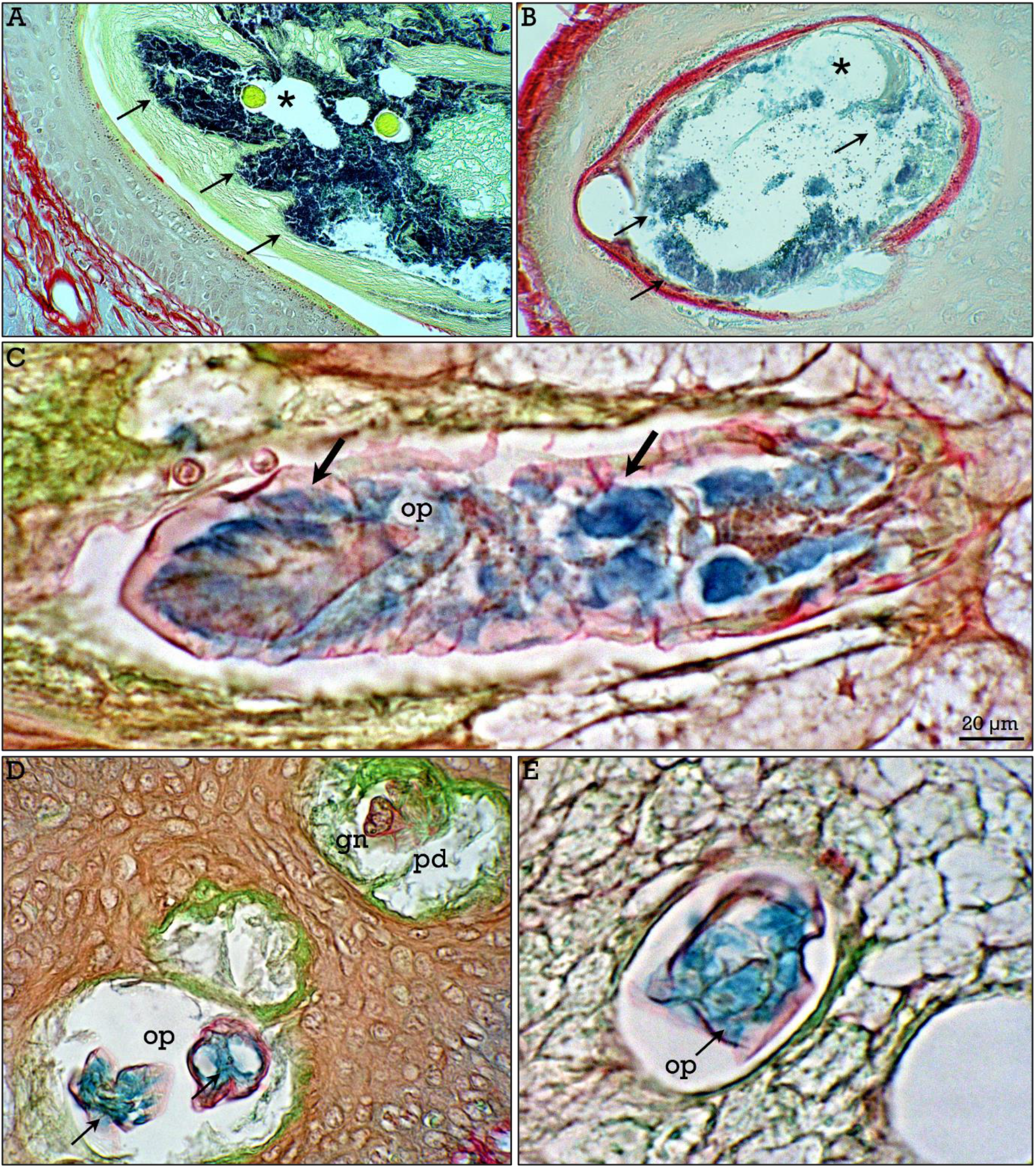
*Relationships between demodex mites and bacteria*. In skin samples showing abundant bacterial colonies (***asterisks***) in the hair follicle infundibula (**A,B**), demodex mites show abundant blue-stained material inside the opistosoma (***arrows*** in **C-F**).

Fungi are not considered as parasites in the strict sense, although some of them have a parasitic lifestyle, and are included as parasites by some authors^**49**^.

Fungi could act as oportunistic infectious agents^**50**^, and several types were evident even in healthy tissue samples, whereas other were clearly involved in pathological processes. Anyway, fungi were clearly stained with RGB trichrome. Some examples from different fungi (*Tinea* spp., *Blastomyces dermatitidis, Cryptococcus neoformans, Aspergillus* spp., and a fungus-like bacteria, *Actinomyces israelii*) are shown in Figure 14, revealing the outstanding staining of these fugi species by RGB trichrome.

**Figure 14.**
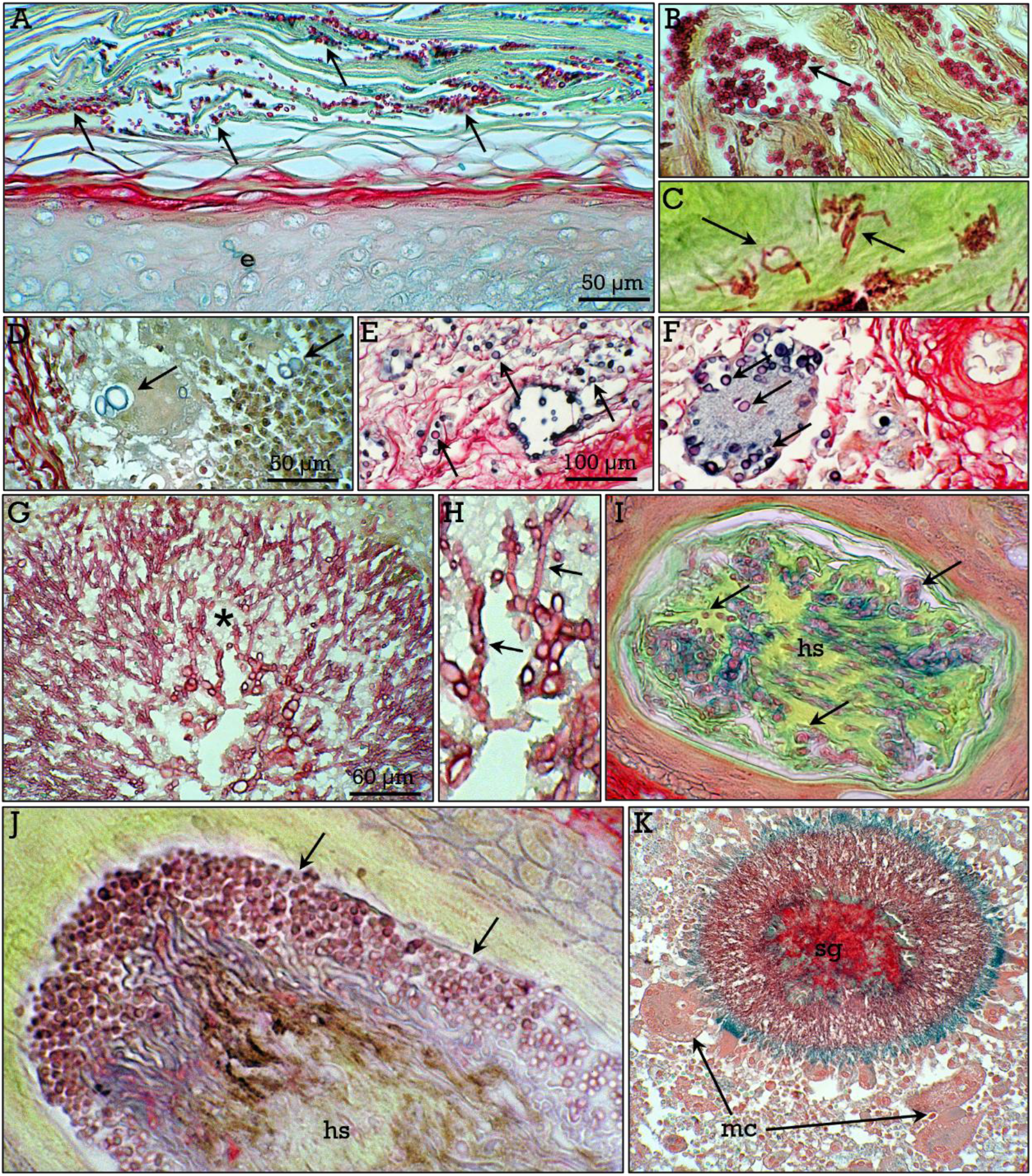
*RGB staining of fungi*. Fungi (***arrows***) stained with RGB trichrome: yeasts (**A,B**) and hyphae (**C**) of fungi in the stratum corneum of normal skin, *Blastomyces* (**D**) and *Aspergillum* (**G,H**) in skin lesions, *Cryptococcus* (**E,F**) in the spleen, and *Tinea* (**I,J**) in hair shafts (***hs***). A filamentous fungus-like (**K**) bacteria (*Actinomyces*) showing a characteristic sulphur granule (***sg***) surrounded by multinucleated cells (***mc***).

Summarizing, the RGB staining allows detailed observation of the structural characteristics of parasites and, at the same time, provides information about host tissue alterations, such as inflammatory reactions at the host-parasite interface and structural changes such as parasite-induced fibrosis, representing a simple and non-expensive histopathological staining method, making it a useful tool for the study of tissues infected by a wide array of pathogens.

## Acknowledgements

The authors are very grateful to María Cantillo and Esteban Tarradas for their technical assistance.

## Author contribution

Both authors were involved in conducting the study and contributed to the final version of the manuscript.

## Declaration of interest

The author declare no competing interests.

## Notes

### Competing Interest Statement

The authors have declared no competing interest.

## References

1. Alum A, Rubino JR, Ijaz MK, et al. The global war against intestinal parasites-should we use a holistic approach? Int J Infect Dis 2010; 14:e732–8.

2. Havelaar AH, Kirk MD, Torgerson PR, et al. World Health Organization Global Estimates and Regional Comparisons of the Burden of Foodborne Disease in 2010. PLoS Med 2015; 12:e1001923.

3. Pisarski K. The Global Burden of Disease of Zoonotic Parasitic Diseases: Top 5 Contenders for Priority Consideration. Trop Med Infect Dis 2019; 4:44.

4. Kantzanou M, Karalexi MA, Vrioni G, et al. Prevalence of Intestinal Parasitic Infections among Children in Europe over the Last Five Years. Trop Med Infect Dis 2021;6:160.

5. Feasey N, Wansbrough-Jones M, Mabey DCW, et al. Neglected tropical diseases. Br Med Bull 2010; 93:179–200.

6. Pullan R, Brooker S. The health impact of polyparasitism in humans: are we underestimating the burden of parasitic diseases? Parasitology 2008; 135:783–94.

7. Clark NJ, Owada K, Ruberanziza E. Parasite associations predict infection risk: incorporating co-infections in predictive models for neglected tropical diseases. Parasites & Vectors 2020; 13:138.

8. Windsor DA. Most of the species on Earth are parasites. Int J Parasitol 1998; 28:1939–41.

9. Dobson A, Lafferty KD, Kuris AM. Colloquium paper: hoimage to Linnaeus: how many parasites? how many hosts? Proc Natl Acad Sci U S A 2008; 105:11482–9.

10. Costello MJ. Parasite rates of discovery, global species richness and host specificity. Integr Comp Biol 2016;56:588–99.

11. Duarte Rocha CF, Bergallo HG, Bittencourt EB. More than just invisible inhabitants: parasites are important but neglected components of the biodiversity. Zoologia 2016; 33:e20150198.

12. Simner PJ. Medical Parasitology Taxonomy Update: January 2012 to December 2015. J Clin Microbiol 2016; 55:43–7.

13. Hemalatha S, Edith R, Sreekumar C. Histopathological detection of parasitic infections. J Vet Parasitol 2018; 32:28–34.

14. Ricciardi A, Ndao M. Diagnosis of parasitic infections: what’s going on? J Biomol Screen 2015; 20:6–21.

15. Momcilovic S, Cantacessi C, Arsic-Arsenijevic V, et al. Rapid diagnosis of parasitic diseases: current scenario and future needs. Clin Microbiol Infect 2019; 25:290–309.

16. Wood JC, Friedly G, de la Maza LM. Detection of helminth ova and larvae in trichrome-stained stool smears. J Clin Microbiol 1982; 16:1137–44.

17. Woods GL, Walker DH. Detection of infection or infectious agents by use of cytologic and histologic stains. Clin Microbiol Rev 1996; 9:382–404.

18. Gaytán F, Morales C, Reymundo C et al. A novel RGB-trichrome staining method for routine histological analysis of musculoskeletal tissues. Sci Rep 2020; 10:16659.

19. Dubey JP, lindsay DS, Speer CA. Structures of Toxoplasma gondii tachyzoites, bradyzoites, and sporozoytes and biology and development of tissue cysts. Clin Microbiol Rev 1998; 11:267–99.

20. Basyoni MMA, Rizk EMA. Nematodes ultrastructure: complex systems and processes. J Parasit Dis 2016; 40:1130–40.

21. Corsi AK, Wightman B, Chalfie M. A Transparent Window into Biology: A Primer on Caenorhabditis elegans. Genetics 2015; 200:387–407.

22. Gottstein B, Pozio E, Nöckler K. Epidemiology, diagnosis, treatment, and control of trichinellosis. Clin Microbiol Rev 2009; 22:127–145.

23. Wendt S, Trawinski H, Schubert S, et al. The diagnosis and treatment of pinworm infection. Dtsch Arztbl Int 2019; 116:213–9.

24. Simón F, Diosdado A, Sles-Lucas M, et al. Human dirofilariosis in the 21st century: A scoping review of clinical cases reported in the literature. Transbound Emerg Dis 2021: doi: 10.1111/tbed.14210

25. Wu Z, Sofronic-Milosavljevic Lj, Nagano I, et al. Trichinella spiralis: nurse cell formation with emphasis on analogy to muscle cell repair. Parasites & Vectors 2008; 1:27.

25. Beiting DP, Park PW, Appleton JA. Synthesis of syndecan-1 by skeletal muscle cells is an early response to infection with Trichinella spiralis but is not essential for nurse cell development. Infect Immun 2006; 74:1941–3.

27. Street JM, Souza ACP, Alvarez-Prats A, et al. Automated quantification of renal fibrosis with Sirius Red and polarization contrast microscopy. Physiol Rep 2014; 2:e12088.

28. Martínez-Criado Y, Millán-López A, Galán N, et al. Apendicitis aguda por Enterobius vermicularis, una etiologia inusual en niños. Rev Esp Enf Dig 2012; 104:396–7.

29. Taghipour A, Olfatifar M, Javanmard M, et al. The neglected role of Enterobius vermicularis in appendicitis: A systematic review and meta-analysis. PLoS One 2020; 15:e0232143.

30. Engin O, Yildirim M, Yakan S, et al. Can fruit seeds and undigested plant residuals cause acute appendicitis. Asian Pac J Trop Biomed 2011;1:99–11.

31. Campora M, Antonelli CT, Grillo F, et al. Seeds in the appendix: a “fruitfull” exploration. Histopathology 2017; 71:322–5.

32. Razzano D, Gonzalez RS. Disease, drugs, or dinner? Food histology can mimic drugs and parasites in the gastrointestinal tract. Virchows Arch 2020; 477:593–5.

33. Grillo F, Campora M, Cornara L, et al. The Seeds of Doubt: Finding Seeds in Intriguing Places. Front Med 2021; 8:655113.

34. Khamesipour F, Nezaratizade S, Basirpour B, et al. review of Dirofilaria spp. infection in human and animals in Iran. Res Vet Sci Med 2021; 1:1.

35. Djakovic I, Lenicek T, Beck R, et al. Subcutaneous Dirofilariasis in Female Pubic Region - Case Report. Open Access Maced J Med Sci 2019; 7:392–5.

36. Pupic-Bakrac A, Pupic-Bakrac J, Beck A, et al. Dirofilaria repens microfilaremia in humans: Case description and literature review. One Healthy 2021; 13:100206.

37. Huang L, Appleton J. Eosinophils in Helminth Infection: Defenders and Dupes. Trends Parasitol 2016; 32:798–807.

38. Budke CM, Carabin H, Ndimubanzi PC. A systematic review of the literature on cystic echinococcosis frequency worldwide and its associated clinical manifestations. Am J Trop Med Hyg 2013; 88:1011–27.

39. Vuitton DA, McManus DP, Rogan MT. International consensus on terminology to be used in the field of echinococcoses. Parasite 2020; 27:41.

40. Thompson RCA. The taxonomy, phylogeny and transmission of echinococcus. Exp Parasitol 2008; 119:439–46.

41. Craig P, Mastin A, van Kesteren F, et al. Echinococcus granulosus: epidemiology and state-of-the-art of diagnostics in animals. Vet Parasitol 2015; 213:132–148.

42. Manterola C, Rojas C, Totomoch-Serra A, et al. Genotipos de Echinococcus granulosus en hidatidosis humana alrededor del mundo. Revision sitemática. Rev Chilena Infectol 2020; 37:541–9.

43. Lacey N, Raaghallaigh SN, Powell FC. Demodex mites--commensals, parasites or mutualistic organisms?. Dermatology 2011; 222: 128–30.

44. Litwin D, Chen W, Dzika E, et al. Human permanent ectoparasites; recent advances on biology and clinical significance of Demodex mites: Narrative review article. Iran J Parasitol 2017; 12:12–21.

45. Merad Y Derrar H, Hebri ST. Demodex, an eclectic mite living in both hair and skin: A review. J Aller Res 2019; 1:1–7.

46. Chen W, Plewig G. Human demodicosis: revisit and a proposed classification. British J Dermatol 2014; 170:1219–25.

47. Kim HS. Microbiota in rosacea. Am J Clin Dermatol 2020; 21:525–35.

48. Namazi MR. A possible role for human follicle mites in skin’s defense against bacteria. Indian J Dermatol Venereol Leprol 2007;73:270.

49. Garcia LS. Classification and nomenclature of human parasites. In: Feigin RD, Cherry J, Demmler-Harrison GJ, Kaplan SL. Feigin and Cherry textbook of pediatric infectious diseases. Philadelphia: Saunders/Elsevier 2009:2861–66.

50. Köhler JR, Casadevall A, Perfect J. The spectrum of fungi that infects humans. Cold Spring Harb Perspec Med 2014; 5:a019273.

